# USP11 promotes endothelial apoptosis-resistance in pulmonary hypertension by deubiquitinating HINT3

**DOI:** 10.1101/2022.05.17.492159

**Authors:** Andrew J. Jang, Victor Tseng, Jae Sun Kim, Robert S. Stearman, Yutong Zhao, Jing Zhao, Jiwoong Choi, John Lister, Michael J. Passineau, Wilbur A. Lam, Changwon Park, Raymond J. Benza, Bum-Yong Kang

## Abstract

**Introduction:** Pulmonary arterial hypertension (PAH) is a progressive, lethal, and incurable disease of the pulmonary vasculature. Evolving evidence indicates that the ubiquitin-specific proteases (USPs), play an important role in the pathogenesis of PAH by deubiquitinating key proteins involved in cell proliferation, migration, and apoptosis. Our genome-wide association study (GWAS) analysis-paired with transcriptomic profiling indicated that deubiquitinase USP11 and histidine triad nucleotide binding protein 3 (HINT3) are positively correlated and that their expression increased in lungs of PAH patients compared to control (fail donor) group, and inversely correlated with survival. However, mechanisms and function of the USP11/HNT3 axis have not been explored in PAH. Therefore, we aimed to investigate that HINT3 stabilized by USP11 activation links to endothelial apoptosis-resistance in PAH.

**Methods and Results:** Expression of USP11 and HINT3 was increased in the lungs of idiopathic PAH (IPAH) patients and Hypoxia/Sugen-treated mice using qRT-PCR and Western blot analyses. USP11 and HINT3 interacted physically as shown by co-immunoprecipitation (co-IP) assay in human pulmonary artery endothelial cells (HPAECs). HINT3 levels were decreased upon transfection of HA-tagged Ubi plasmid into HPAECs. Pretreatment with the potent proteasome inhibitor MG132 prolonged the half-life of HINT3 protein, indicating that HINT3 is degraded by polyubiquitination. HINT3 was stabilized and destabilized by forced overexpression or siRNA knockdown of USP11 respectively. Similarly, treatment with mitoxantrone, a USP11 antagonist, reduced HPAEC HINT3 expression. HINT3 interacted with the antiapoptotic mediator, BCL2. Overexpression of USP11 increased BCL2 content, congruent to elevated lung tissue levels seen in IPAH patients and Hypoxia/Sugen-treated mice. Conversely, knockdown of HINT3 function led to depletion of BCL2.

**Conclusions:** The HINT3-USP11 axis contributes to apoptosis-resistance in pulmonary artery endothelial cells, as is potentially a novel and attractive therapeutic target for ubiquitination modulators.

## INTRODUCTION

Pulmonary arterial hypertension (PAH) is a progressive and incurable disease marked by pulmonary vascular occlusive remodeling leading to eventual right ventricular failure and death.^1-3^ Even though effective therapy has improved survival rate, the effective 5-year survival is only ∼60%, and females account for 79.5% in all prevalent cases amongst those with PH enrolled in the registry to evaluate early and long-term PAH disease management (REVEAL registry).^4-6^

PH is characterized by progressive increase in pulmonary vascular resistance due to endothelial dysfunction, abnormal proliferation of pulmonary vascular wall cells, and vascular remodeling. Evolving evidence indicate that the ubiquitin-specific proteases (USPs), deubiquitinating enzymes play important roles in cell proliferation, migration, and apoptosis by removing ubiquitin groups from unanchored polyubiquitin chains in the pathogenesis of PH.^7-10^ USPs regulate ubiquitination-meditated degradation and stabilize proteins to maintain the response to cellular level signals and altered environmental conditions. This stabilization is established when the deubiquitinating enzymes remove ubiquitin from ubiquitin-conjugated protein substrates to regulate the stability of a target protein. For example, ubiquitin-specific proteases 11 (USP11) which belongs to the USP family has been known as a pro-inflammatory mediator and regulates apoptosis, proliferation, cancer chemoresistance, ferroptosis, autophagy.^11-14^ However, little is known about the role and mechanisms of USP11 in PAH pathogenesis.

Recent studies demonstrated that PAH lung transcriptome analysis can provide expression patterns specific to PAH subtypes, clinical parameters, and lung pathology variables.^15^ Built on these datasets, we identified associations between survival and single nucleotide polymorphisms (SNPs) in or near genes thought to be relevant to PAH pathogenesis. One of these SNPs, we discovered that histidine triad nucleotide binding protein 3 (HINT3) shows a strong association with worsened survival in PAH. HINT3 is a nucleotide hydrolase and transferase. However, the expression, function, and disease-specific action of HINT3 is still unknown. Further, we found ubiquitin specific peptidase 11 (USP11) and HINT3 are strongly associated with PAH. However, the mechanisms and function of USP11/HNT3 axis are poorly understood in PH. Therefore, we hypothesized that HINT3 stabilized by USP11 activation links directly to endothelial anti-apoptosis in PH.

During PAH pathogenesis, pulmonary artery endothelial cells become hyperproliferative and resistant to apoptosis, leading to neointimal thickening and narrowing of the vessel lumen.^16-18^ Apoptosis is a cell death program which prevents uncontrolled cell replication. For example, B-cell lymphoma 2 (BCL2) is a gene that once expressed promotes cell survival by blocking apoptosis.^19^ BCL2 is located in the membranes of the mitochondria, the endoplasmic reticulum membrane and the nuclear envelope. BCL2 in the mitochondria prevents the release of apoptogenic factors, which leads to the occurrence of mitochondrial dysfunction.^20, 21^ However, mechanisms if HINT3 regulates BCL2 expression in PH are not defined. Further, we have observed increases in BCL2 levels in endothelial cells harvested from catheter tips of patients with PAH,^22^ suggesting that BCL2 might contribute directly to regulation of anti-apoptosis in PAH. Therefore, we further hypothesized that USP11/HINT3 axis regulates the activity and the abundance of the BCL2 proteins, which could lead to cell proliferation of pulmonary vessels in PH.

In this study, we explored the role and interaction of USP11 and HINT3 in PH. We show that levels of USP11 and HINT3 are increased in the lungs of IPAH patients and Hypoxia/Sugen-treated mice. Further, we demonstrate that USP11 enhances the stability of HINT3 by deubiquitinating HINT3 and thereby increases anti-apoptotic marker BCL2 levels. Our findings suggest that inhibition of USP11/HINT3 axis may act as a novel therapeutic target in PH pathogenesis.

## MATERIALS AND METHODS

### Control fail donor and IPAH lung tissues

We obtained de-identified peripheral lung tissues from control or idiopathic pulmonary arterial hypertension (IPAH) patient specimens collected by the Pulmonary Hypertension Breakthrough Initiative (PHBI). IPAH samples were derived from 4 male and 5 female patients, 29-55 years old whereas control specimens were derived from 2 male and 2 female subjects, 23-56 years old who were failed donors.

### Reagents

Human pulmonary artery endothelial cells (HPAECs) were obtained from Cell Applications, Inc. (San Diego, CA, USA), GAPDH, and β-actin (ACTB) antibodies from Cell Signaling (Danvers, MA, USA). USP11, HINT3, and BCL2 antibodies were purchased from Abcam (Waltham, MA, USA), fetal bovine serum (FBS), dimethyl sulfoxide (DMSO), from Invitrogen (Waltham, MA, USA) and USP11 inhibitor, mitoxantrone from Millipore Sigma (Burlington, MA, USA).

### *In vivo* mouse model of PH

Male C57BL/6J mice, aged 8-12 weeks, were treated three times with weekly injections of the VEGF receptor antagonist, Sugen 5416 (SU, 20 mg/kg, subcutaneous injection) and exposed to hypoxia (HYP/SU, 10% O_2_) or normoxia (NOR/SU, 21% O_2_) for 3 weeks as reported previously.^23^ All animals were given unrestricted access to water and standard mouse chow. All animal studies were approved by the Institutional Animal Care and Use Committee of Emory University or the Atlanta Veterans Affairs Healthcare System.

### *In vitro* human pulmonary artery endothelial cells

HPAECs were cultured in Endothelial Cell Growth Media (Cell Applications, San Diego, CA, USA) containing 1% (*vol*/*vol*) penicillin/streptomycin. Cells were grown under standard incubator condition at 37°C and with 5% CO_2_. HPAECs cells were plated in 6-well plate or 10 cm cell culture dishes. Media was changed every 3 days and cells were split when the cell density reached 80-90% confluence.

### Gain or loss of USP11 or HINT3 function in HPAECs

For loss of USP11 or HINT3 function (LOF), HPAECs were transfected with scrambled or USP11 (F:5’-GUCAAUGAGAAUCAGAUCGAGUCCA-3’, R:3’-GACAGUUACUCUUAGUCUAGCUCAGGU-5’) or HINT3 (F:5’-CUUGCAGUGAUAUAAUUGACAACAT-3’, R:3’-UUGAACGUCACUAUAUUAACUGUUGUA-5’) dicer-substrate siRNA (DsiRNA) (20 nM, Integrated DNA Technologies, Coralville, IA) using Lipofectamine 3000 transfection reagent (Invitrogen) according to the manufacturer’s instructions. After transfection for 6 hours, the transfection media were replaced with EGM containing 5% FBS and incubated at room temperature for 72 hours. In selected experiments, HPAECs were treated with the USP11 inhibitor, mitoxantrone for 0-4 h. HPAEC lysates were then harvested and examined for USP11, HINT3, and BCL21 levels using Western blot assays. To overexpress USP11 (gain of function), HPAECs were transfected with USP11 plasmid constructs (1 μg, oxUSP11) or empty vector (Mock) we previously reported.^23^ After transfection for 6 hours, media were replaced with fresh 5% FBS EGM, and HPAEC lysates were then harvested and examined for USP11, HIN3, and BCL2 levels using Western blot assays.

### Co-immunoprecipitation

To ascertain the presence of protein-protein interaction between USP11-HINT3 or HINT3-BCL2, we performed co-immunoprecipitation (co-IP) with HINT3 antibody assay in HPAECs. Briefly, for co-IP, about 1 mg cell lysates were incubated with specific primary antibody overnight and were then added to 40 μL protein A/G agarose beads (Thermo Fisher Scientific, catalog 78609) for an additional 2 hours at room temperature. The precipitation complex was washed by cold PBS-T (PBS+ 0.1% Tween 20) three times for 5 minutes each. The precipitation was added to 50 μL lysis buffer and analyzed by immunoblotting with an enhanced chemiluminescence detection system (Advansta, catalog K-12043-D20). Equal loading of protein samples from each group was evaluated using β-actin or GAPDH antibody after using Restore WB stripping buffer (Thermo Fisher Scientific, catalog 21059).

### HINT3 half-life and ubiquitination

To more characterize if HINT3 protein is degraded by ubiquitination, HPAECs were transfected with either control plasmids (empty vector) or 1μg of HA-ubiquitin plasmids (HA-tagged Ubi) using Lipofectamine 3000 (Thermo fisher Scientific, PA, USA) according to the manufacturer’s recommendations. Twenty-four hours later, HPAEC lysates were collected in 1X M-PERTM Mammalian Protein Extraction Reagent from Thermo Scientific (Waltham, MA, USA) spiked with protease/phosphatase inhibitors examined by immunoblotting. ACTB levels were used as an endogenous control. To examine half-life of USP11, HPAECs were cultured in complete endothelial cell growth media (EGM) containing 1% (vol/vol) penicillin/streptomycin. Twenty-four hours after plating, HPAECs were exposed to 20 μg/mL cycloheximide (CHX) in the presence or absence of 20 μM MG132 for 0, 2, 4, or 8 h.

### mRNA quantitative real-time polymerase chain reaction (qRT-PCR) analysis

To measure USP11, HINT3, and BCL2 levels in the lungs of IPAH patients or mouse lungs, total RNAs were isolated using the total RNA isolation kit (Qiagen, USA). Human: USP11, forward: 5’-TCCTCAGCCCAGAGTGTTCT-3’, reverse: 5’-CACACACACAGCAGAAGGTACA-3’, HINT3, forward: 5’-AGTGCGTTAAGTTCCCTGATG-3’, reverse: 5’-CCACCCACTCTCAGAAGTGTT-3’, BCL2, forward: 5’-GTGACCGTACAGTCGGGATT-3’, reverse: 5’-GCCGTACAGTTCCACAAAGG-3’ and mouse: Usp11, forward: 5’-CACCCTCTCCTGGTGTCAGT-3’, reverse: 3’-ATCTGAGGTGGGTTTGGTCA-5’), Hint3 forward: 5’-TGACAGCAACTGCGTGTTCT-3’, reverse :3’-CCGCTGGTTTGATATCTTTG-5’), and Bcl2, forward: 5’-GTACCTGAACCGGCATCTG-3’, reverse:3’-GCTGAGCAGGGTCTTCAGAG-5’) mRNA levels mRNA levels in the same sample were determined and quantified using specific mRNA primers. Human GAPDH, forward: 5’-GCCCAATACGACCAAATCC-3’, reverse: 3’-AGCCACATCGCTCAGACAC-5’) and mouse Gapdh, forward: 5’-AGCTTGTCATCAACGGGAAG-3’, reverse: 3’-TTTGATGTTAGTGGGGTCTCG-5’) mRNA levels were used as an endogenous control.

### Western blot analysis

All protein homogenates from human and mouse lungs or HPAECs were subjected to Western blot analysis as reported.^23^ Primary antibodies were purchased from Abcam (Waltham, MA, USA) and included USP11 anti-rabbit polyclonal antibody (1:250 dilution, Cat #ab109232, 110 kDa), HINT3 Rabbit polyclonal antibody (1:500 dilution, Cat # ab121960, 20 kDa), BCL2 Rabbit polyclonal antibody (1:500 dilution, Cat #ab59348, 26 kDa). GAPDH rabbit polyclonal antibody (1:10,000 dilution, Cat # G9545, 37 kDa) or ACTB Rabbit polyclonal antibody (1:10,000 dilution, Cat #4970, 37 kDa) was purchased from Cell signaling (Danvers, MA, USA). Proteins were visualized using infrared secondary antibodies (1:10,000) using Rad gel doc proprietary software. Relative protein levels were visualized using image Lab software (Bio-Rad, Hercules, CA, USA), quantified Image J software, and normalized to GAPDH or ACTB levels within the same lane.

### Statistical Analysis

For all measurements, data were presented as mean ± standard error of the mean (SE). All data were analyzed using analysis of variance (ANOVA). Post hoc analysis used the Student Neuman Keuls test to detect differences between specific groups. In studies comparing only two experimental groups, data were analyzed with Student’s t-test to determine the significance of treatment effects. Statistical significance was defined as p< 0.05. Statistical analyses were performed using GraphPad Prism, Version 9.0 software (LaJolla, CA, USA).

## RESULTS

### USP11 and HINT3 are upregulated in the lung tissue from PAH patients and from Hypoxia/Sugen-treated mice *in vivo*

We revisited our recent genome-wide association study (GWAS) analysis using Affymetrix microarray^15^ and found that USP11 and HINT3 are positively correlated, and their expression increased in lungs of PAH patients compared to failed donor controls (**Supplementary Figure S1**). Therefore, we validated the findings by measuring levels of USP11 and HINT3 mRNAs and proteins in the lung tissues from idiopathic PAH (IPAH) patients and Hypoxia/Sugen-treated mice. USP11 mRNA (**Figures 1A** and **1C**) and protein (**Figures 1B** and **1D**), and concordantly, HINT mRNA (**Figures 2A** and **2C**) and protein (**Figures 2B** and **2D**), were significantly increased in the lungs of IPAH patients and Hypoxia/Sugen-treated mice *in vivo*.

**Figure 1.**
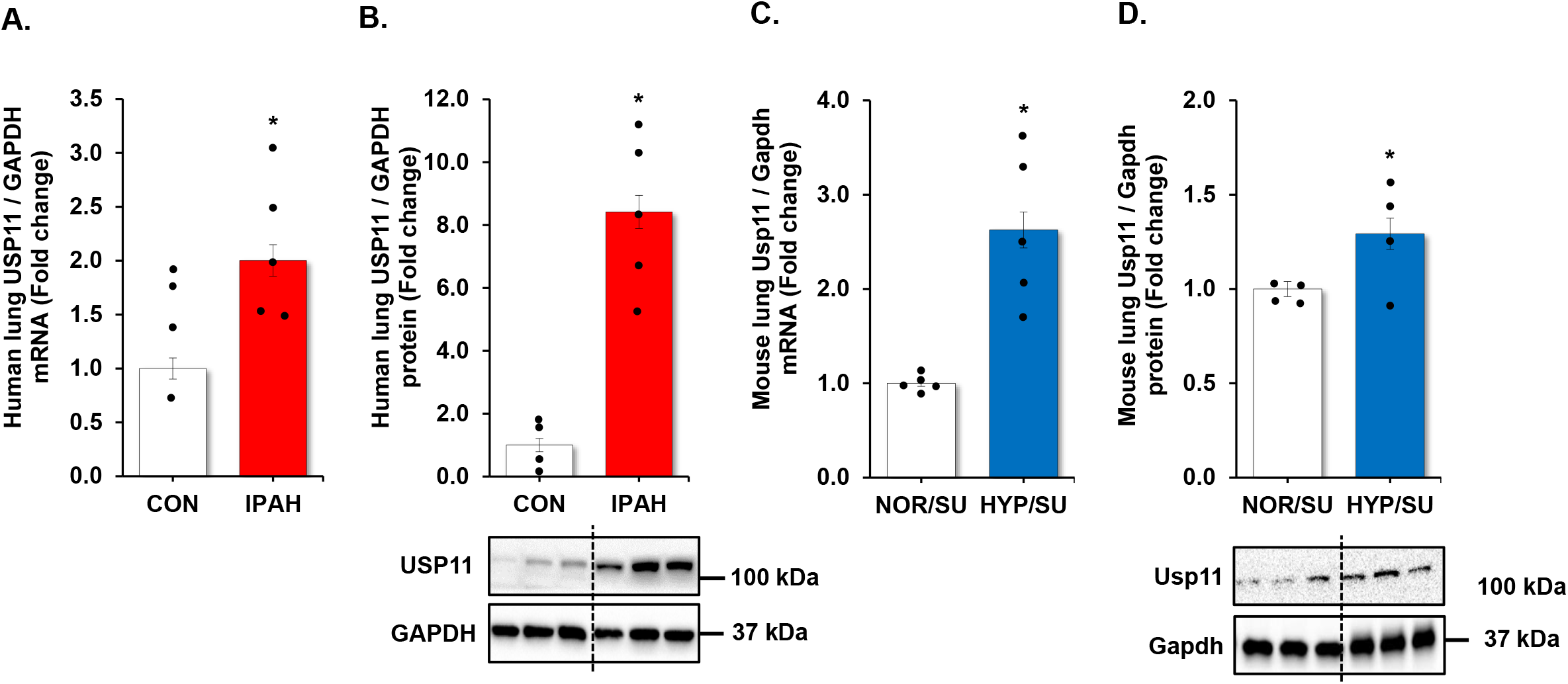
UPS11 is increased in clinical and experimental PH samples. **(A and B)** fail donor (CON) or idiopathic pulmonary arterial hypertension (IPAH) patient lungs were subjected to qRT-PCR and Western blots. n=4-5. **(C and D)** Whole lung homogenates were collected from normoxic/sugen (NOR/SU) or hypoxic/sugen (HYP/SU)-treated mice. All bars represent mean USP11 mRNA or protein levels ± SE relative to GAPDH expressed as fold-change vs. control (CON) or vs. NOR/SU. n=4-5. *p<0.05 vs respective control condition.

**Figure 2.**
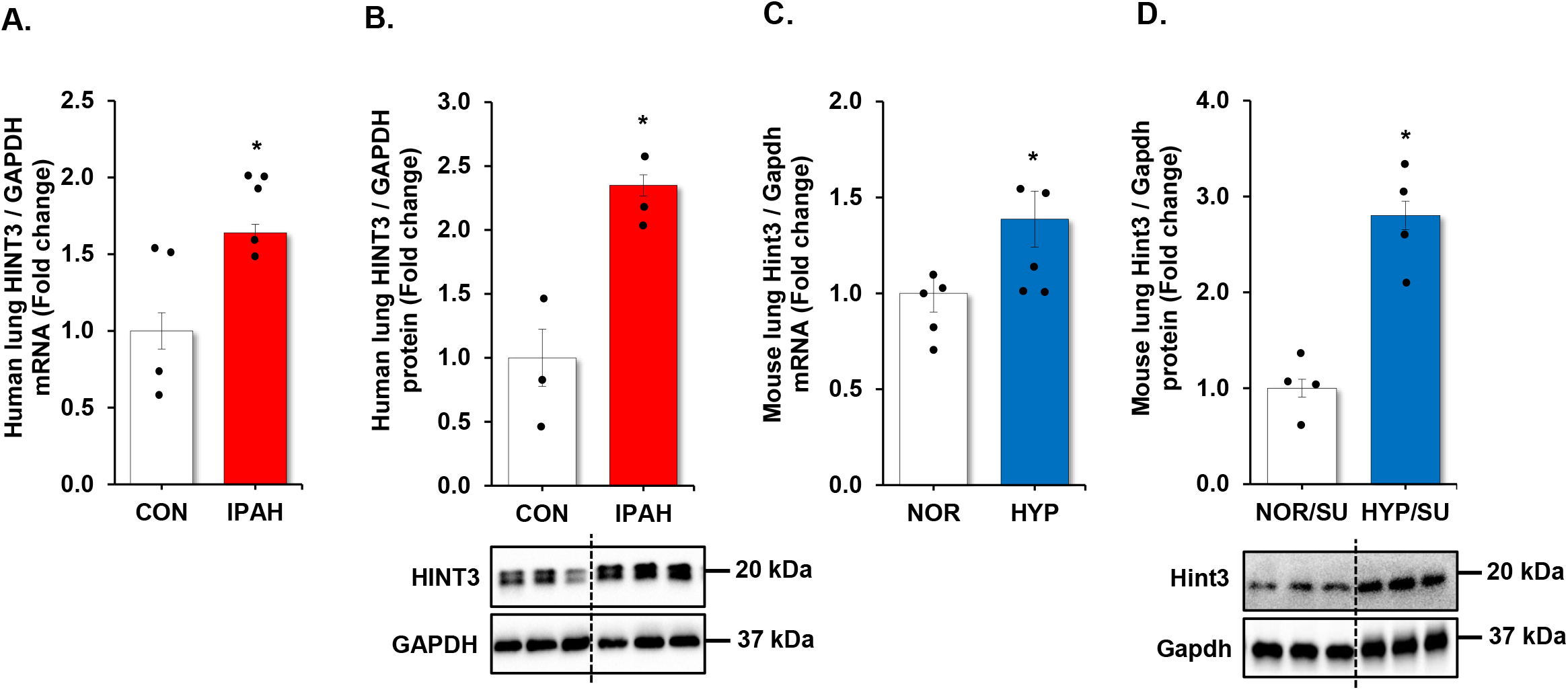
HINT3 is increased in clinical and experimental PH samples. **(A and B)** fail donor (CON) or idiopathic pulmonary arterial hypertension (IPAH) patient lungs were subjected to qRT-PCR and Western blots. n=4-5. **(C and D)** Whole lung homogenates were collected from normoxic/sugen (NOR/SU) or hypoxic/sugen (HYP/SU)-treated mice. All bars represent mean HINT3 mRNA or protein levels ± SE relative to GAPDH expressed as fold-change vs. control (CON) or vs. NOR/SU. n=4-5. *p<0.05 vs respective control condition.

### HINT3 is degraded by ubiquitination

To understand the functional significance of the association of USP11 and HINT3, we sought to protein-protein interactions between USP11 and HINT3 with. co-immunoprecipitation (co-IP) assay in HPAECs. As illustrated in **Figure 3A**, HINT3 strongly binds USP11. Since USP11 has deubiquitination ability, we hypothesized that HINT3 is degraded by ubiquitination. To address the hypothesis, first HPAECs were transfected with either control (empty vector) or 1μg of HA-tagged ubiquitin plasmid (HA-Ubi), and the level of HINT3 was measured. As shown in Figure 3B, HINT3 protein was reduced upon transfection with HA-Ubi i, indicating -mediated degradation of HINT3.-. To further dissect mechanisms of HINT3 protein degradation, HPAECs were treated in a time-dependent manner with the translational inhibitor, cycloheximide (CHX), to arrest *de novo* protein synthesis. As illustrated in **Figure 3C**, CHX treatment revealed a half-life of HINT3 of 8 hours. To determine whether HINT3 degradation was primarily dependent on the ubiquitin proteasome pathway, HPAECs were pretreated with the proteasome inhibitor, MG132, followed by CHX. MG132 significantly prolonged the half-life of HINT3 protein, with only 10% degradation at 8 hours. These data suggest that ubiquitination is a major regulatory step governing steady-state HINT3 levels.

**Figure 3.**
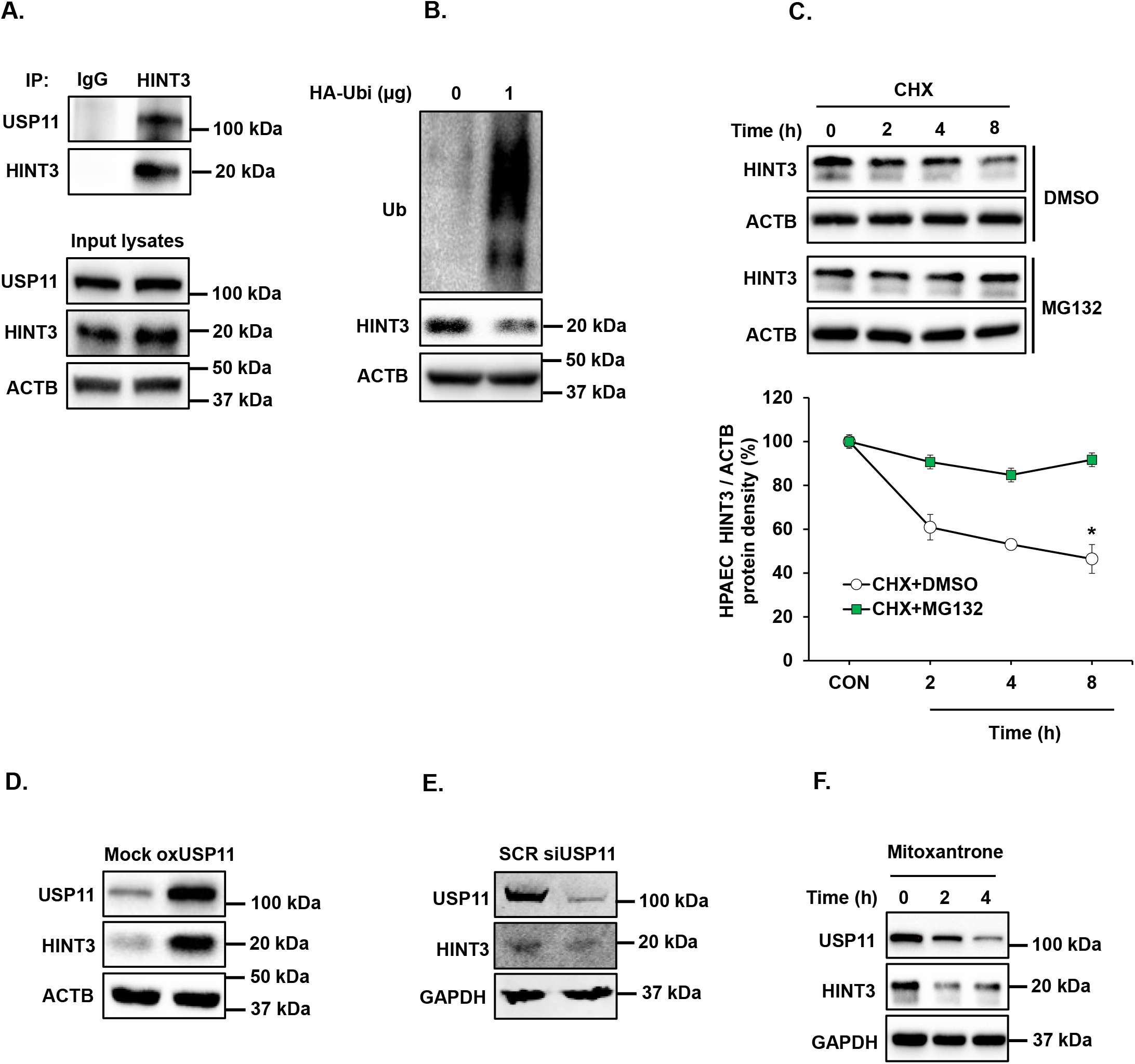
HINT3 is degraded by ubiquitination and USP11 deubiquitinates HINT3 in HPAECs *in vitro*. **(A)** Human pulmonary artery endothelial cells (HPAECs) were cultured for 48 h. Cell lysates were harvested and subjected to immunoprecipitation (IP) with either anti-IgG antibody or anti-HINT3 antibody, and then immunoblotted with anti-USP11, anti-HINT3, or anti-β-actin (ACTB) antibody. **(B)** HPAECs were transfected with either the control plasmid (empty vector) or HA-ubiquitin (1 μg, HA-Ubi) plasmid for 24 h. HPAECs were collected and assayed for anti-HA, anti-HINT3, and ACTB (loading control) antibody by immunoblotting. **(C)** HPAECs were pretreated with DMSO or MG132 for 2 h and were treated with cycloheximide (CHX, 20 μg/ml) for 0, 2, 4, and 8 h to inhibit de novo protein synthesis and harvested for Western blotting. The levels of HINT3 at time 0 was set as 100% and the percent HINT3 protein remaining following CHX treatment at each time point was calculated accordingly. All bars represent mean HINT3 protein levels ± SE relative to ACTB expressed as fold-change vs. CON *p<0.05 vs CON, n=3. **(D)** HPAECs were transfected with either the control plasmid (Mock) or USP11 (1 μg, oxHUWE1) plasmid for 6 h, media were replaced with endothelial growth medium (EGM) containing 5% FBS. And then incubated for an additional 72 h. Cell lysates were collected and immunoblotted with anti-HINT3 or ACTB antibody. **(E)** HPAECs were treated with scrambled (SCR) or HINT3 (20 nM) siRNAs for 6 h, media were replaced with endothelial growth medium (EGM) containing 5% FBS. And then incubated for an additional 72 h. Western blotting was performed for HINT3 or GAPDH protein. **(F)** HPAECs) were treated with dimethyl sulfoxide (DMSO) or USP11 inhibitor (Mitoxantrone) for 0-4 h. Western blotting was performed for HINT3 or GAPDH protein. n=3.

### USP11 deubiquitinates HINT3 in HPAECs *in vitro*

Having established that USP11 interacts with HINT3, and that HINT3 degradation is regulated by ubiquitination, we tested whether USP11 acts as a deubiquitinase to stabilize HINT3. First, HPAECs were transfected with USP11 plasmid or empty vector (mock) for 72 hours. Overexpression of USP11 increased HINT3 levels (**Figure 3D**). However, knockdown of USP11 with siRNA reduced HINT3 expression in HPAECs (**Figure 3E**). We further explored whether pharmacologic inhibition of USP11 using mitoxantrone affected HINT3 expression. As shown in **Figure 3F**, mitoxantrone significantly reduced USP11 and HINT3 expression. Taken together, these findings suggest that USP11 interacts with HINT3 to stabilize through deubiquitination.

### USP11 regulates the anti-apoptotic mediator BCL2 via HINT3 binding activity

To further clarify the relationship between USP11, HINT3, and BCL2, we first examined BCL2 expression using qRT-PCR and Western blot analysis. Congruent with previous results,^24, 25^ lung tissue levels of BCL2 were increased in IPAH patients (**Figures 4A** and **4B**) and Hypoxia/Sugen-treated mice (**Figures 4C** and **4D**). Next, we performed co-IP to determine whether BCL2 interacts with HINT3 and found the interaction between BCL2 and HINT3 (Figure 5A) Interestingly, siRNA knockdown of HINT3 attenuated BCL2 expression in HPAECs, while USP11 levels remained unchanged (**Figure 5B**). Conversely, overexpression of USP11, which can indirectly increase the expression of HINT3, significantly augmented expression ofBCL2 (**Figure 5C**). These findings suggest that USP11 deubiquitinates HINT3, allowing it to positively regulate BCL2 expression.

**Figure 4.**
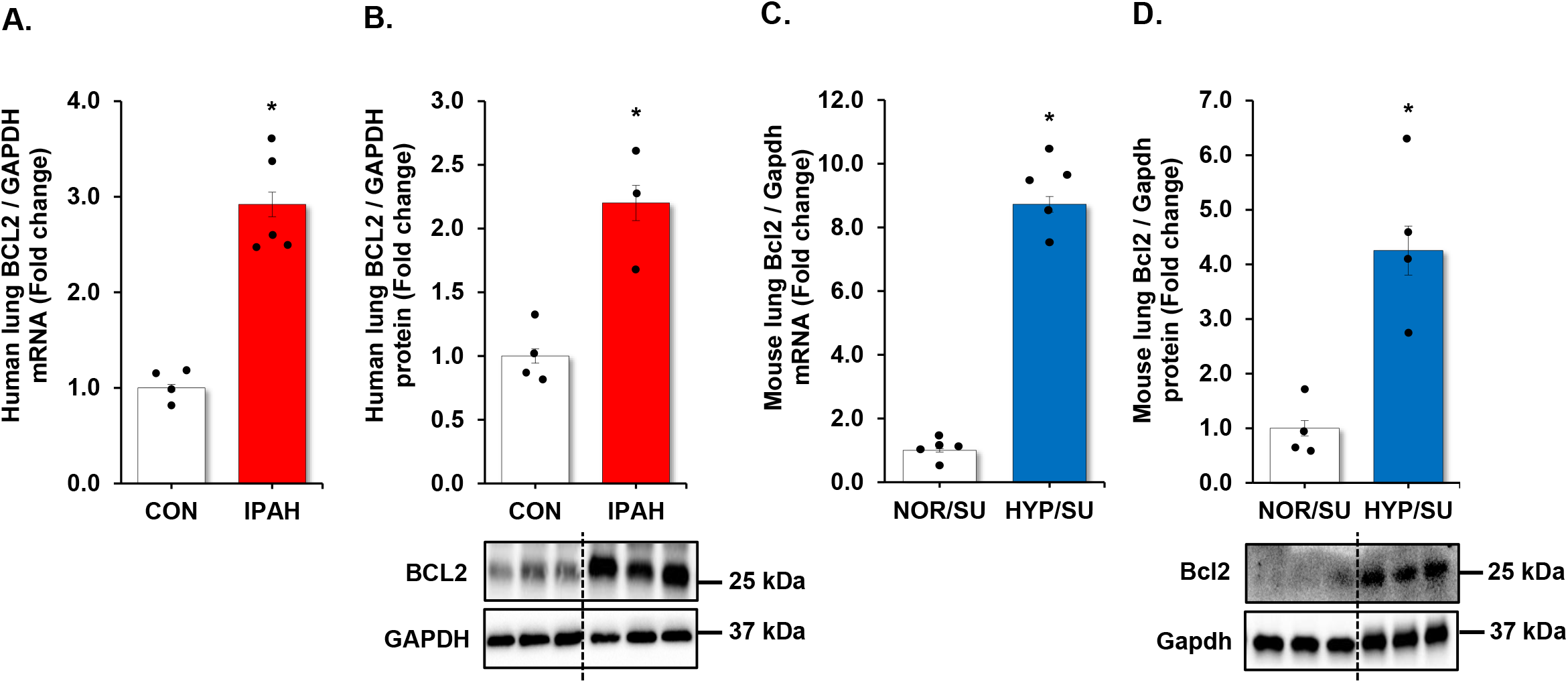
BCL2 is increased in clinical and experimental PH samples. **(A and B)** fail donor (CON) or idiopathic pulmonary arterial hypertension (IPAH) patient lungs were subjected to qRT-PCR and Western blots. n=4-5. **(C and D)** Whole lung homogenates were collected from normoxic/sugen (NOR/SU) or hypoxic/sugen (HYP/SU)-treated mice. All bars represent mean BCL2 mRNA or protein levels ± SE relative to GAPDH expressed as fold-change vs. control (CON) or vs. NOR/SU. n=4-5. *p<0.05 vs respective control condition.

**Figure 5.**
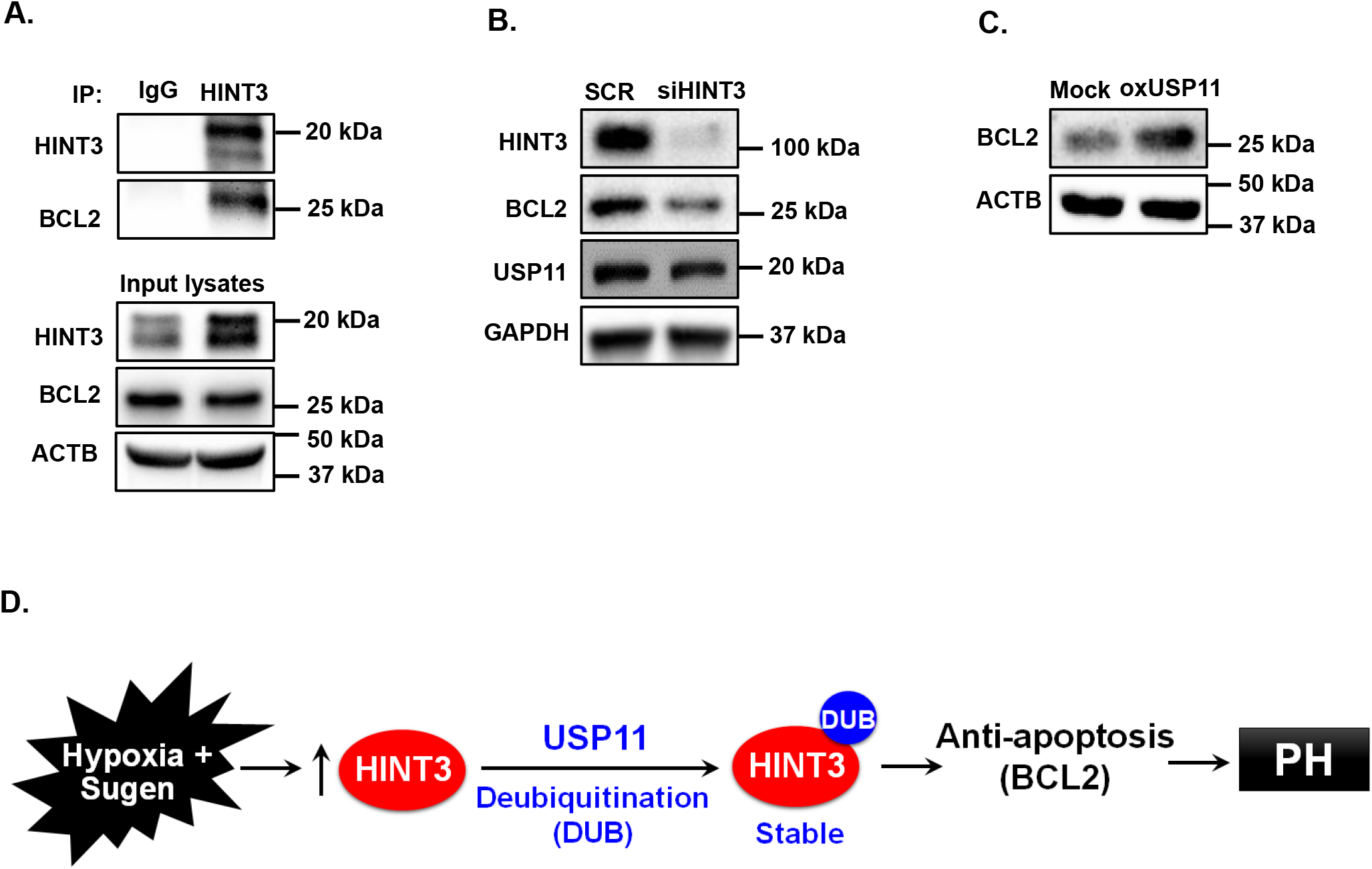
HINT3 regulates BCL2 expression in HPAECs *in vitro*. Human pulmonary artery endothelial cells (HPAECs) were cultured for 48 h. Cell lysates were harvested and subjected to immunoprecipitation (IP) with either anti-IgG antibody or anti-HINT3 antibody, and then immunoblotted with anti-HINT3, anti-BCL2, or anti-β-actin (ACTB) antibody. **(B)** HPAECs were treated with scrambled (SCR) or HINT3 (20 nM) siRNAs for 6 h, media were replaced with endothelial growth medium (EGM) containing 5% FBS. And then incubated for an additional 72 h. Western blotting was performed for HINT3, BCL2, USP11, or GAPDH protein. **(C)** HPAECs were transfected with either the control plasmid (Mock) or USP11 (1 μg, oxUSP11) plasmid for 6 h, media were replaced with endothelial growth medium (EGM) containing 5% FBS. And then incubated for an additional 72 h. Cell lysates were collected and immunoblotted with anti-BCL2 or ACTB antibody. **(D)** Hypothetical schema defining the role of USP11/HINT3/BCL2 signaling in PH pathogenesis. Hypoxia induces USP11 that stabilizes HINT3 levels by HINT3 deubiquitination. Increases in HINT3 stimulate anti-apoptosis marker, BCL2 expression promoting PH pathogenesis.

## DISCUSSION

This study provides new evidence of close mechanistic interplay between novel candidate genes identified on our recent genome-wide assoociation study of pulmonary arterial hypertension (PAH). Specifically, polymophisms in the debiquitinating enzyme USP11, and the adenosine 5’-monophosphoramidase HINT3, were previously unreported in PAH, but found to to be prognostic for survival. In this work, we sought to understand the regulatory relationships between USP11 and HINT3, and establish a link to apoptosis-resistane, a pathologic phenotype acquired in pulmonary vascular wall cells in PAH.

Our findings can be summarized as follows: (1) both USP11 and HINT3 are regulated at multiple levels, based on concordant elevation of mRNA and protein in the lungs of IPAH patients and Hypoxia/Sugen-treated mice with experimental PAH *in vivo*; (2) however, the balance of ubiquitination and USP11-mediated deubiquitination is a major determinant of HINT3 protein stability; and (3) HINT3 positively regulates the antiapoptotic mediator BCL2, consistent with the stereotyped finding of apoptosis-resistance in pulmonary artery endothelial cells. Immunoprecipitation assays corroborate interactions between relevant binding partners: USP11-HINT3--BCL2. Therefore, abnormal upregulation of USP11 contributes to hypo-ubiquitination and inappropriate stabilization of HINT3, resulting in downstream activation of BCL2.

The homeostasis of many cellular proteins is regulated by ubiquitin-mediated proteostasis in response to environmental stimuli. Ubiquitination is a prolific post-translational modification that regulates diverse processes by branding proteins for degradation either by the proteasome or lysosome.^26^ Various ubiquitin ligases have been described in neoplastic and degenerative disease processes.^27^ Conversely, removal of ubiquitin chains from ubiquitinated proteins is catalyzed by deubiquitinating enzymes (DUBs), which can rescue substrate proteins from degradation^26^. Abnormal ubiquitination of several vasoactive, redox, and mitogenic proteins such as calveolin-1, angiotensin converting enzyme-2, superoxide dismutase-2, followed by increased proteosomal degradation, have been reported in PAH.^28-30^

Recently, our group demonstrated a global change in the hypoxic lung ‘ubiquitome’, with differential ubiquitination detected in at least 131 proteins.^31^ Across the lysine landscape, multiple sites of the proteins were hypo-ubiquitinated, which may also indicate increased DUB activity stimulated by hypoxia. Therefore, the contribution of DUBs to PAH pathogenesis warrants more careful scrutiny. USP11 is a newly described DUB whose repertoire of targets is unknown. Based on the strength of association between the pair in our GAWAS dataset, we hypothesized that HINT3 could be a downstream target of USP11.

HINT3 is a poorly characterized protein with unknown function other than AMP hydrolase activity of uncertain significance. There is a paucity of literature on the relevance of HINT3 to disease, but existing reports indicate high expression in breast cancer, where it strongly correlates with mortality,^32^ and in hepatocellular carcinoma, where it associated with apoptosis-resistance in the face of serum starvation.^33^ These findings indicate that HINT3 may potentially function as a proto-oncogene. Pursuant to this notion, we demonstrated that HINT3 directly interacts with and positively regulates the anti-apoptotic protein BCL2. Notably, we have discovered that a series of polymorphisms in non-coding regions of HINT3, resulting in its upregulation in PAH lungs, are associated with poor survival in PAH (data reserved for separate publication). These findings represent additional novel insights into the clinical significance of HINT3 in PAH, a condition which features cancer-like expansion of pulmonary vascular wall cells.

The current study has several important limitations. To confirm the contribution of USP11-HINT3 dysfunction to PAH, pulmonary artery hemodynamics, right venticular hypertrophy, and lung vascular remodeling must be directly measured in the endothelial-targeted USP11 overexpressing trangenic mouse model. To solidify the impact of HINT3 on pro-remodeling phenotypes, additonal markers of apoptosis, endothelial dysfunction. proliferation, contractile-secretory trandifferentiation, and migration should be assessed. Cell-type specificity of this pathway, encompassing pulmonary artery smooth muscle cells and adventitial fibroblasts, should similarly be examined. Additionally, to demonstrate the therapeutic feasibility of targeting the USP11-HINT3 axis, studies are needed to investigate whether pharmacologic inhibition or genetic ablation of USP11 attenuates rodent models of PAH *in vivo*.

In summary, the current study mechanically validates the relationship between two novel co-regulated targets detected on genomic and transcriptomic screening. We demonstrate that HINT3 is stabilized through UPS11-directed deubiquitination. Stabilized HINT3 upregulation enhanced BCL2 activation in HPAECs. We speculate that mitoxantrone, which is an FDA-approved antineoplastic agents with canonical DNA topoisomerase inhibitor activity, may be repurposed or functionalized to target USP11 to restore apoptosis-sensitivity and reduced vascular remodeling in PAH.^34^ Furthermore, our results suggest that HINT3 polymorphisms may be prospectively validated in longitudinal patient cohorts as a mechanistic biomarker in PAH.

## AUTHORSHIP

Conception, hypothesis delineation, and design, A.J.J. and B.Y.K.; acquisition of data, analysis, and interpretation, A.J.J., C.P., J.S.K., V.T., R.S.S., Y.Z., J.Z., J.C., J.L., R.J.B., M.J.P., W.A.L., and B.Y.K.; writing of the article, A.J.J., C.P., J.S.K., V.T., J.C., J.L., W.A.L., and B.Y.K.

## Acknowledgements

This study was supported by funding from NIH National Heart, Lung, and Blood Institute R01 grants (HL119291 to CP, HL157164 to YZ, HL151513 to JZ, and HL133053 to BYK). Data/Tissue samples provided by PHBI under the Pulmonary Hypertension Breakthrough Initiative (PHBI). Funding for the PHBI is provided under an NHLBI R24 grant, #R24HL123767, and by the Cardiovascular Medical Research and Education Fund (CMREF).

## DISCLOSURES

The contents do not represent the views of the Department of Veterans Affairs or the United States Government.

## Notes

### Competing Interest Statement

The authors have declared no competing interest.

