## Supplemental Figures for "USP11 promotes endothelial apoptosis-resistance in pulmonary hypertension by deubiquitinating HINT3"

#### SUPPLEMENTARY FIGURE LEGENDS

**Supplementary Figure. S1. PAH exhibits upregulated USP11, HINT3, and BCL2 expression.**

RNA sequencing was conducted using PAH human vs failed donor normal lung. Data are expressed as mean USP11, HINT3, or BCL2 ± SE

### SUPPLEMENTARY FIGURE LEGENDS

#### USP11 promotes endothelial apoptosis-resistance in pulmonary hypertension by deubiquitinating HINT3

Andrew J. Jang, PhD[1]*, Victor Tseng[2], Jae Sun Kim[3], Robert S. Stearman, PhD[4], Yutong Zhao, MD PhD[5], Jing Zhao, MD PhD[5], Jiwoong Choi[6], John Lister[1], Michael J. Passineau, PhD[1], Wilbur A. Lam[7,11], Changwon Park, PhD[8], Raymond J. Benza, MD[9], and Bum-Yong Kang, PhD[3,10,11]*

[1]Division of Hematology and Cellular Therapy, Allegheny Health Network Cancer Institute, Pittsburgh, PA, USA.
[2]Respiratory Medicine, Ansible Health; Mountain View, California, USA.
[3]Department of Medicine, Division of Pulmonary, Allergy, Critical Care, and Sleep Medicine, Emory University School of Medicine, Atlanta, GA, USA.
[4]Department of Medicine, University of Indiana, Indianapolis, Indiana, USA.
[5]Department of Physiology and Cell Biology, Department of Internal Medicine, The Ohio State University, Columbus, OH, USA
[6]Division of Pulmonary, Critical Care, and Sleep Medicine, The University of Kansas School of Medicine, Kansas City, KS. USA
[7]Georgia Institute of Technology, Atlanta, GA, USA.
[8]Department of Cellular and Molecular Physiology, Louisiana State University Health Science Center, Shreveport, LA, USA.
[9]The Ohio State University Wexner Medical Center, Columbus, Ohio, USA.
[10]Atlanta Veterans Healthcare System, Decatur, GA, USA.
[11]Department of Pediatrics, Division of Hematology, Oncology, and BMT, Emory University School of Medicine, Atlanta, GA, USA.

### Supplementary Fig. S1

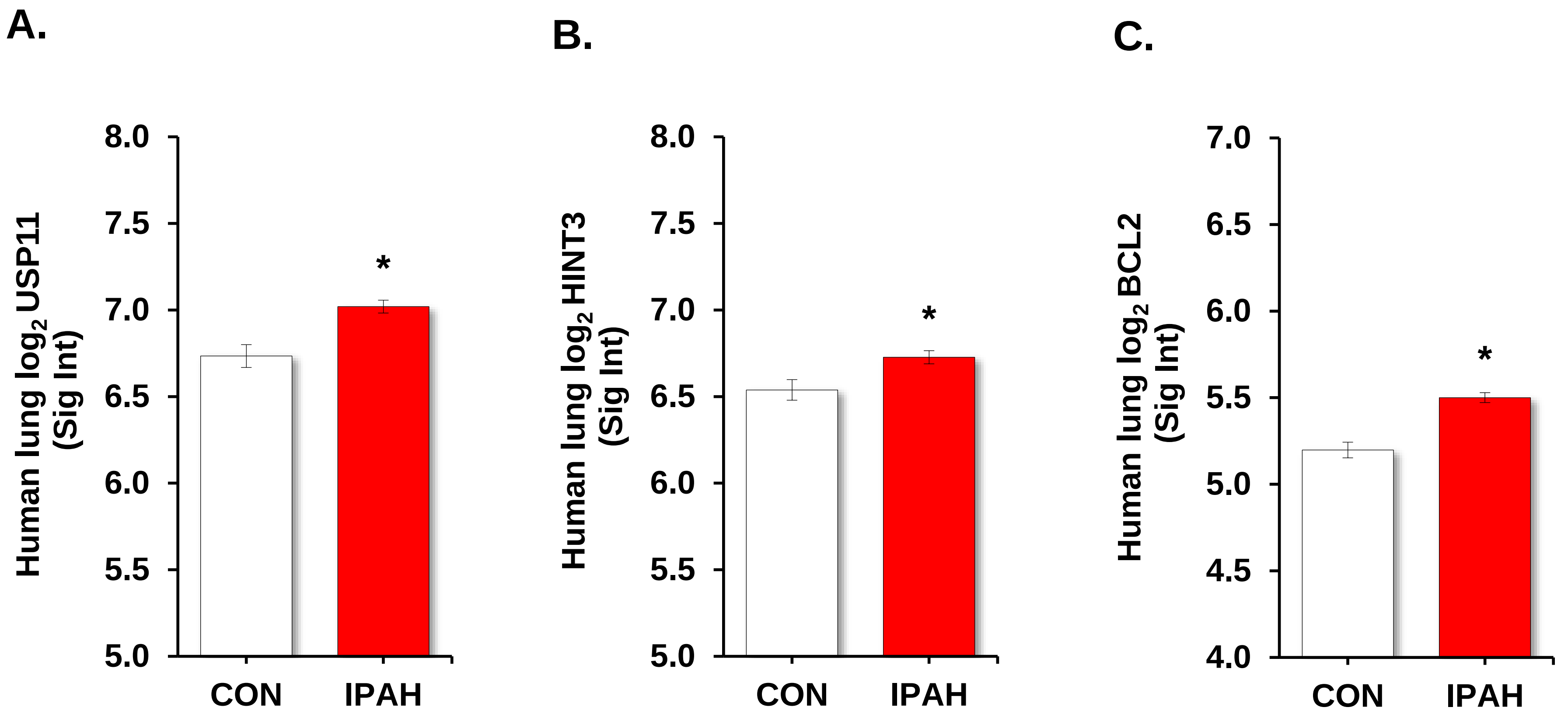
